# CCR5 promotes the migration of CD8^+^ T cells to the leishmanial lesions

**DOI:** 10.1101/2023.10.10.561700

**Authors:** Laís Amorim Sacramento, Camila Farias Amorim, Claudia G. Lombana, Daniel Beiting, Fernanda Novais, Lucas P. Carvalho, Edgar M. Carvalho, Phillip Scott

## Abstract

Cytolytic CD8^+^ T cells mediate immunopathology in cutaneous leishmaniasis without controlling parasites. Here, we identify factors involved in CD8^+^ T cell migration to the lesion that could be targeted to ameliorate disease severity. CCR5 was the most highly expressed chemokine receptor in patient lesions, and the high expression of CCL3 and CCL4, CCR5 ligands, was associated with delayed healing of lesions. To test the requirement for CCR5, *Leishmania-*infected Rag1^-/-^ mice were reconstituted with CCR5^-/-^ CD8^+^ T cells. We found that these mice developed smaller lesions accompanied by a reduction in CD8^+^ T cell numbers compared to controls. We confirmed these findings by showing that the inhibition of CCR5 with maraviroc, a selective inhibitor of CCR5, reduced lesion development without affecting the parasite burden. Together, these results reveal that CD8^+^ T cells migrate to leishmanial lesions in a CCR5-dependent manner and that blocking CCR5 prevents CD8^+^ T cell-mediated pathology.

## INTRODUCTION

Cutaneous leishmaniasis is caused by an intracellular protozoan parasite transmitted by sandflies. The disease exhibits a broad spectrum of clinical manifestations ranging from self-healing lesions to extensive mucosal damage. While some patients resolve their lesions spontaneously, others develop lesions that progress to chronicity and lead to the development of severe mucosal disease. Importantly, the severe disease in many patients is due to the inflammatory response rather than uncontrolled parasite replication [1–5]. Thus, treatment of cutaneous leishmaniasis may require not only anti-parasitic drugs but also host-directed therapies to limit inflammation.

Cytolytic CD8^+^ T cells play a pathological role in cutaneous leishmaniasis and contribute to the chronicity of the disease. Experimental murine models and transcriptional studies in patients’ lesions demonstrate that cytolytic CD8^+^ T cells mediate increased pathology by promoting extensive cytolysis, leading to inflammasome activation and interleukin-1β (IL-1β) release, which in turn feeds the inflammation and enhances the magnitude of the disease [6–10]. In previous studies, we found that inhibition of cytotoxicity [6] or the subsequent events, including inflammasome activation and IL-1β release, blocked severe disease [10,11]. Because CD8^+^ T cells were implicated in treatment failure in *L. braziliensis* patients in previous studies[7], here we investigate the factors that govern the recruitment of those cells in order to identify additional therapeutic approaches to improve the treatment of the disease.

We performed transcriptional analysis of lesions from *L. braziliensis* patients and experimental models to address the factors involved in CD8^+^ T cell migration to cutaneous leishmaniasis lesions. We identified *CCR5* as the most highly expressed chemokine receptor in *L. braziliensis*-lesions. Moreover, patients with a high expression of the CCR5 ligands, *CCL3* and *CCL4*, exhibited delayed healing of lesions and had elevated expression of key cytolytic genes. Additionally, we found that IL-15 upregulates CCR5 expression on CD8^+^ T cells in patients. Translating the findings to the murine model, we found that CD8^+^ T cells express CCR5 preferentially at the lesions and that deletion of CCR5 in CD8^+^ T cells dampens pathology in a model of severe lesions induced by *L. braziliensis*. Importantly, mice treated with maraviroc (MVC), a selective inhibitor of CCR5, significantly reduced lesion development without affecting parasite number. Collectively, these results show that CCR5 mediates the migration of CD8^+^ T cells to leishmanial lesions and identify a possible target for host-directed therapy by using pharmacological inhibition of CCR5 to prevent CD8^+^ T-cell mediated pathology.

## RESULTS

### Genes encoding for CCR5 and its ligands are overexpressed in patient lesions and are associated with delayed healing

We previously demonstrated that cytotoxic CD8^+^ T cells mediate the development of severe lesions in cutaneous leishmaniasis [7,8]. In order to characterize what drives the migration of CD8^+^ T cells to lesions, we created a list of chemokine receptors associated with T cell migration based on the literature [12–19]. We took advantage of our published human RNA-seq dataset of lesions from *L. braziliensis* patients [7] and evaluated the expression of chemokine receptors and their ligands in biopsies from patients’ lesions compared to healthy skin (Fig 1A). The analysis showed that among the genes investigated, *CCR5* was the most highly expressed in lesions, and *CCL3* and *CCL4*, which bind CCR5, were also enriched in lesions (Fig 1B). *CCL3* and *CCL4* are statistically correlated in *L. braziliensis*-lesions (*\*\*p = 0.0049*), so we stratified the cohort of patients in half (normal distributions) based on the high (CCL3/4^high^) or low (CCL3/4^low^) expression of these genes (Fig 1C) and used this strategy to evaluate the impact on the healing time.

**Fig 1.**
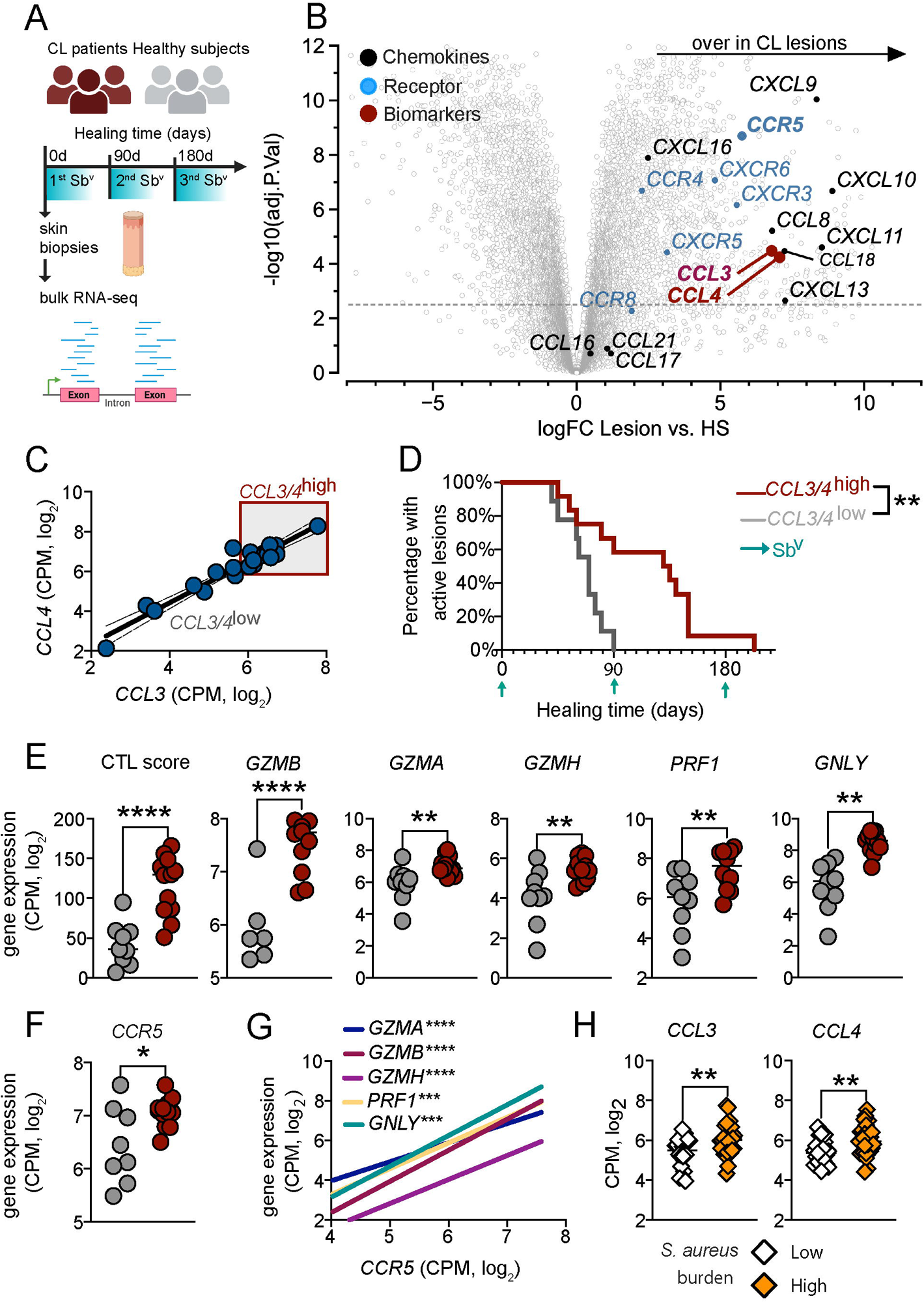
*CCR5* and its ligands are overexpressed in patient lesions and are associated with delayed healing. (A) The study design for the bulk RNA-seq dataset. Lesions were collected at day 0, and the complete re-epithelialization of lesions was followed at days 90 and 180 of antimony treatment. RNAseq analysis of skin or lesions was performed with 7 healthy subjects (HS) and 21 cutaneous leishmaniasis patients (CL). (B) Volcano plot highlighting overexpressed chemokines (black), chemokines receptors (blue), and biomarkers that predict treatment outcome (red) in biopsies from patients relative to biopsies from healthy subjects. (C) Correlation between *CCL4* and *CCL3* expression. *CCL3/4*^high^ expression was estimated based on *CCL3* > 5.8 and *CCL4* > 6. (D) Healing time of patients with high and low *CCL3/4* expression. (E) MCP counter abundance for cytotoxic T lymphocyte (CTL) score and *GZMB*, *GZMA*, *GZMH*, *PRF1*, and *GNLY* expression between patients with high and low expression of CCL3/4. (F) *CCR5* expression in patients with high and low *CCL3/4* expression. (G) Correlation between *CCR5* expression and *GZMB*, *GZMA*, *GZMH*, *PRF1*, and *GNLY*. (H) *CCL3* and *CCL4* expression in patients with high and low *S. aureus* transcriptional abundance expression. Data was obtained from RNASeq analysis of lesions from 51 patients. Gene expression is represented as counts per million (CPM) in the log2 scale. \**p < 0.05, **p* ≤ *0.01, ***p* ≤ *0.001, ****p < .0001*.

As demonstrated in Fig 1D, patients with CCL3/4^high^ expression exhibited delayed healing compared to CCL3/4^low^ patients. Since the increased pathology observed in *L. braziliensis* patients is mediated by the cytolytic activity of CD8^+^ T cells [8,20,21], we asked if CCL3/4^high^ expression was associated with CD8^+^ T cells. Using MCP-counter as a method to estimate cell abundances from unstructured RNAseq data[22], we found an association between CCL3/4^high^ expression and cytotoxic T lymphocyte abundances (CTL score) (Fig 1E). Given that cytotoxicity-related genes are associated with treatment outcome [7], we further evaluated whether the genes encoding for the cytolytic machinery were associated with CCL3/4^high^ expression. We observed an overexpression of *GZMB*, *GZMA*, *GZMH*, *PRF1,* and *GNLY* in patients with CCL3/4^high^ expression compared to CCL3/4^low^ expression (Fig 1E). Additionally, patients with CCL3/4^high^ expression exhibited increased *CCR5* expression, suggesting that the high abundance of these chemokines leads to the recruitment of CCR5^+^ cells (Fig 1F). Furthermore, we found a significant positive correlation between *CCR5* expression and *GZMB*, *GZMA*, *GZMH*, *PRF1,* and *GNLY* at lesions (Fig 1G). Since the high abundance of *S. aureus* in *L. braziliensis* lesions is associated with increased expression of cytolytic genes and delayed healing [23], we next evaluated if there is an association between the expression of *CCL3* and *CCL4* with a high or low transcriptional abundance of *S. aureus.* We found that patients with a high abundance of *S. aureus* have an overexpression of *CCL3* and *CCL4* (Fig 1H). Together, these data demonstrate that CCR5 and its ligands are enriched in patients’ lesions and are associated with a delayed healing time of lesions.

### CCL3 and CCL4 are enriched systemically in patients and are associated with cytolytic gene expression

Despite being a localized skin infection, the systemic transcriptional signatures of *L. braziliensis* patients reflect pathways that are present in leishmanial lesions [24]. To address whether *CCR5* and its ligands were overexpressed at the systemic level, we analyzed an RNA-seq dataset from the peripheral blood of 50 cutaneous leishmaniasis patients and 14 healthy subjects (Fig 2A). We observed that *CCL3* and *CCL4* were elevated in the peripheral blood of *L. braziliensis* patients relative to healthy subjects (Fig 2B). Additionally, there was a positive correlation between the expression of *CCL3*, *CCL4*, and *CCR5* with the expression of cytolytic genes *GZMB*, *GZMA*, *GZMH*, *GNLY,* and *PRF1* (Fig 2C-E). These data suggest that the systemic transcriptional signature for CCR5 and its ligands observed in cutaneous leishmaniasis patients recapitulates the signature in lesions. Together, combining the transcriptional data from lesions and peripheral blood, our results suggest that CCR5 may be involved in the migration of CD8*^+^* T cells to the site of infection.

**Fig 2.**
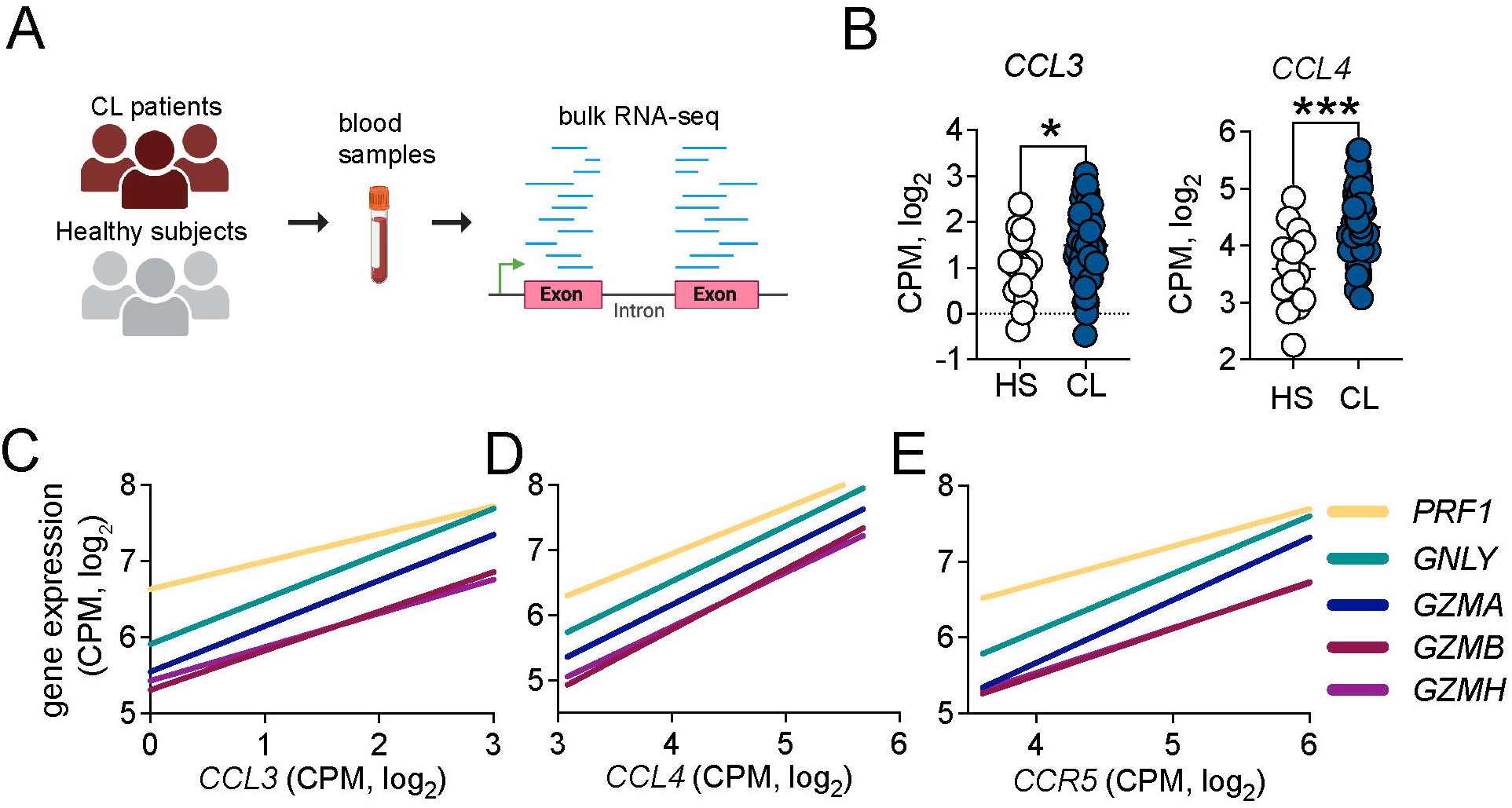
Systemic expression of CCR5 ligands is associated with cytolytic molecules. (A) RNASeq analysis of peripheral blood from 14 healthy subjects (HS) compared to 50 cutaneous leishmaniasis patients (CL). (B) Gene expression of *CCL3* and *CCL4.* (C-E) Correlation of *GZMA*, *GZMB*, *GZMH*, *GLNY*, and *PRF1* with (C) *CCL3*, (D) *CCL4*, and (E) *CCR5*. Gene expression is represented as counts per million (CPM) in the log2 scale. \**p < 0.05, **p* ≤ *0.01, ***p* ≤ *0.001, ****p < .0001*.

### IL-15 upregulates CCR5 expression on CD8^+^ T cells from patients

*IL15* signaling is elevated in *L. braziliensis* lesions[25], and its inhibition lessens CD8*^+^* T cell-mediated pathology[6]. Given the role of IL-15 driving CD8^+^ T cell migration by upregulating CCR5 expression [13], we focused our attention on exploring whether IL15 contributes to the upregulation of CCR5 on CD8^+^ T cells. We first evaluated *IL15* expression by analyzing the RNA-seq dataset from the peripheral blood [24]. We observed that *IL15* is enriched systemically in *L. braziliensis* patients compared to healthy subjects (Fig 3A). Additionally, there was a significant positive correlation between the expression of *IL15* and *CCR5* in the peripheral blood of patients (Fig 3B). No correlation was found in healthy subjects (Fig 3C). These data suggest that the increase in *CCR5* expression is likely because of increased *IL15* expression in patients. To address the direct effect of IL-15 in the upregulation of CCR5 on CD8^+^ T cells from *L. braziliensis* patients, we obtained peripheral blood mononuclear cells (PBMCs) from patients or healthy subjects and stimulated the cells with recombinant IL-15. After 18h, CCR5 expression by CD8^+^ T cells was evaluated by flow cytometry. We observed that IL-15 stimulation enhances the frequency of CD8^+^ T cells expressing CCR5 and the median fluorescence intensity (MFI) from patients (Fig 3D). In contrast, the baseline expression of CCR5 was lower in healthy subjects, and the increase induced by IL-15 was variable (Fig 3E). Collectively, the results demonstrate that IL-15 upregulates CCR5 expression on CD8^+^ T cells from *L. braziliensis* patients.

**Fig 3.**
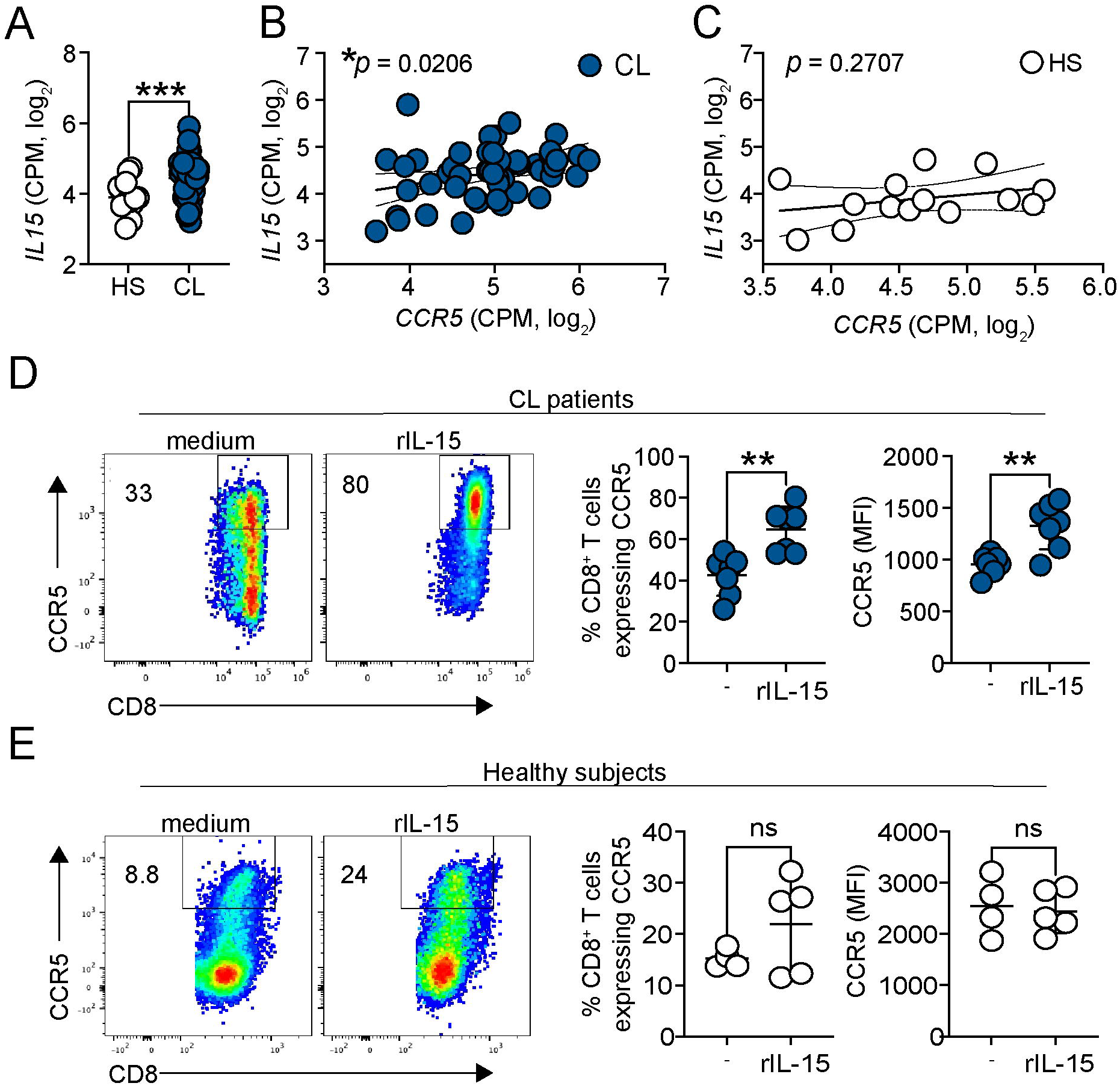
IL-15 upregulates CCR5 expression on CD8^+^ T cells from patients. RNASeq analysis of peripheral blood from 14 healthy subjects (HS) compared to 50 cutaneous leishmaniasis patients (CL). (A) Gene expression of *IL15*. (B and C) Correlation between *IL15* and *CCR5* expressions of (B) CL and (C) HS. Gene expression is represented as counts per million (CPM) in the log2 scale. (D and E) PBMCs from (D) cutaneous leishmaniasis patients and (E) healthy subjects were cultured with recombinant IL-15 for 18h and stained for flow cytometry. Dot plots and graph bars represent the percentage and median fluorescence intensity (MFI) of CCR5 expression by CD8^+^ T cells after IL-15 stimulation. Data were obtained from 5 HS and 7 cutaneous leishmaniasis patients. CL, cutaneous leishmaniasis; HS, healthy subjects; PBMC, peripheral blood mononuclear cells. \**p < 0.05, **p* ≤ *0.01*.

### CD8^+^ T cells express CCR5 in leishmanial lesions in mice

The results of transcriptional analysis of lesions and peripheral blood from patients suggest that CCR5 promotes CD8^+^ T cell migration to lesions. To determine if CCR5 was associated with more severe disease in mice, we evaluated *Ccl3*, *Ccl4,* and *Ccr5* gene expression in lesions from two murine models (Fig 4A and 4C). The skin microbiome exacerbates cutaneous leishmaniasis[23], and colonization of mice with *Staphylococcus xylosus* is associated with more severe lesions [26]. Transcriptional analysis of bulk RNA-seq of infected ears demonstrated that *ccl3*, *ccl4*, and *ccr5* were significantly induced in *S. xylosus*-colonized and *L. major*-infected mice, compared to mice only *S. xylosus*-colonized or *L. major*-infected (Fig 4B).

**Fig 4.**
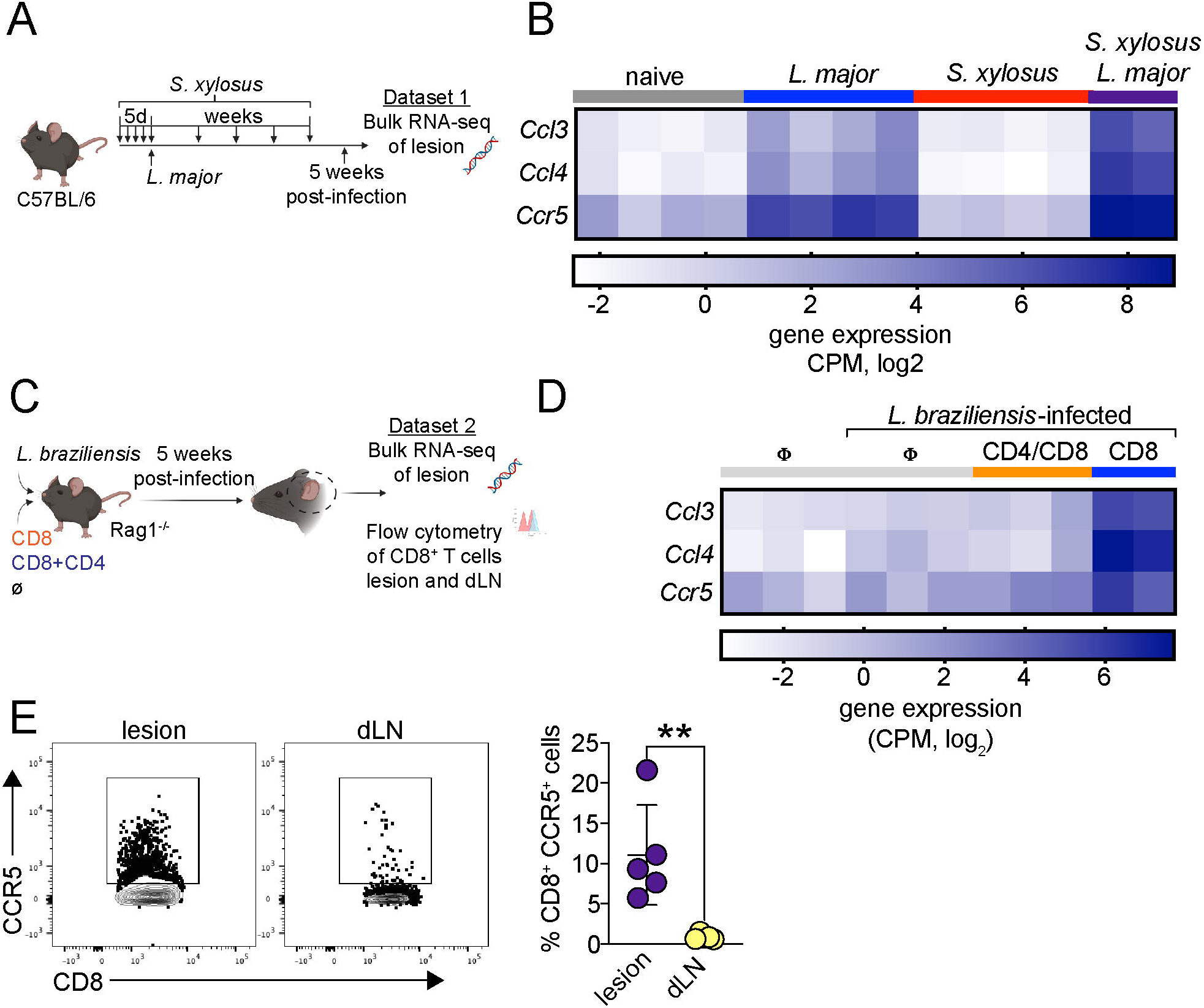
CD8^+^ T cells express CCR5 in leishmanial lesions. (A) C57BL/6 mice were topically colonized with 10^8^ *S. xylosus* every day for a total of five applications and then intradermally infected with 2 x 10^6^ *L. major* parasites. Mice were colonized once per week for the duration of the experiment. At 5 weeks post-infection, lesions were obtained for bulk RNA-seq analysis. (B) Heat map showing gene expression of *Ccl3*, *Ccl4,* and *Ccr5* in the lesions of naïve C57BL/6 mice, *L. major*-infected, *S. xylosus*-colonized or *S. xylosus*-colonized and *L. major*-infected mice (*S. xylosus + L. major*). (C) Rag1^-/-^ mice were infected with *L. braziliensis* in the ear and reconstituted with either CD8^+^ T cells (CD8) or CD8^+^ and CD4^+^ T cells (CD8+CD4) or did not receive cells (Ø). At the peak of infection, lesions were obtained for bulk RNA-seq analysis. (D) Heat map showing gene expression of *Ccl3*, *Ccl4,* and *Ccr5* in the lesions of naïve Rag1^-/-^ mice and infected-Rag1^-/-^ mice not reconstituted with cells and reconstituted with CD4^+^/CD8^+^ or only CD8^+^ cells. Gene expression is represented as counts per million (CPM) in the log2 scale. (E) Representative dot plots and graph bars of CD8^+^ T cells expressing CCR5 in the lesion and dLN of *L. braziliensis*-infected Rag1^-/-^ reconstituted with CD8^+^ T cells. The data (C) are expressed as the means ± SEMs and are representative of two independent experiments (n = 3-5 mice). *\*\*p < 0.01*.

We also examined *Ccl3*, *Ccl4*, and *Ccr5* expression in an experimental model that mimics the pathologic profile of cytotoxic CD8^+^ T cells observed in human lesions [7,8]. In this model, the reconstitution of Rag1^-/-^ mice with CD8^+^ T cells alone leads to the development of uncontrolled lesions in a perforin-dependent manner [8]. The severe pathology is dependent on the cytolytic function of perforin and granzyme B [8] as well as the production of IL-1b [10], similar to what is described in patients’ lesions and is associated with a large increase of neutrophils [7,8]. Transcriptional analysis of infected ears demonstrated that *Ccl3*, *Ccl4*, and *Ccr5* were significantly induced in *L. braziliensis*-infected Rag1^-/-^ lesions that received only CD8^+^ T cells, compared to mice that received both CD4+CD8 cells or mice that did not receive T cells (Fig 4D). CD8^+^ T cells from *L. braziliensis*-infected Rag1^-/-^ mice were also analyzed by flow cytometry, and 10% of CD8^+^ T cells from infected ears expressed CCR5, while less than 1% of CD8^+^ T cells obtained from dLN expressed CCR5 (Fig 4E). These data suggest that CD8^+^ T cells preferentially express CCR5 in the lesion compared to the dLN. Taken together, these results demonstrate that CCR5, as well as its ligands, is highly expressed in the murine models of severe human leishmaniasis.

### CD8^+^ T cell migration to the lesion is dependent on CCR5

To directly test if CCR5 expression on CD8^+^ T cells was required to promote disease, *L. braziliensis*-infected Rag1^-/-^ mice were reconstituted with WT or CCR5^-/-^ CD8^+^ T cells, and the course of infection was monitored (Fig 5A). As expected, Rag1^-/-^ mice reconstituted with WT CD8^+^ T cells (WT CD8) developed uncontrolled lesions. Importantly, Rag1^-/-^ mice reconstituted with CCR5^-/-^ CD8^+^ T cells (CCR5^-/-^ CD8) exhibited significantly smaller lesions (Fig 5B) with less pathology (Fig 5C and 5D). In contrast, parasite burdens were similar in Rag1^-/-^ mice that received WT or CCR5^-/-^ CD8^+^ T cells (Fig 5E).

**Fig 5.**
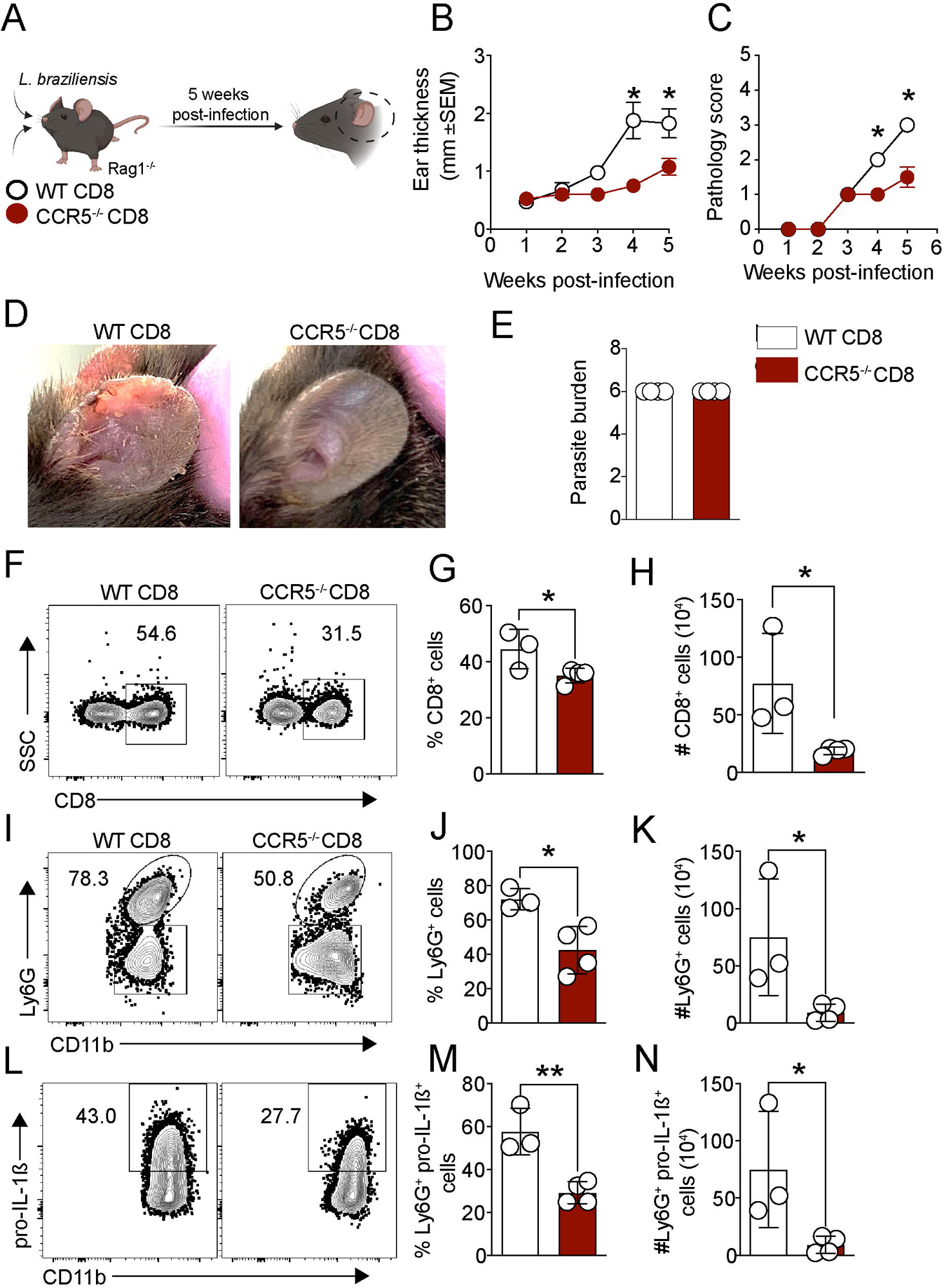
CD8^+^ T cell migration to the lesion is CCR5-dependent. (A) Rag1^-/-^ mice were infected with *L. braziliensis* and reconstituted with WT or CCR5^-/-^ CD8^+^ T cells. (B) Ear thickness and (C) pathology score was assessed weekly. (D) Representative pictures of lesions from infected-Rag1^-/-^ that received WT CD8 or CCR5^-/-^ CD8^+^ T cells at 5 weeks post-infection. (E-N) At 5 weeks post-infection, mice were euthanized, and the ears were digested for (E) parasite quantification by limiting dilution and (F-N) flow cytometry analysis. (F) Representative dot plots and graph bars of the (G) frequency and (H) number of CD8^+^ T cells. (I-N) The neutrophil number (CD11b^+^ Ly6G^+^ cells) and pro-IL-1β expression in the lesion were determined directly *ex vivo* by flow cytometry. (I-K) Representative dot plots and graph bars of neutrophils and (L-N) pro-IL-1β expression. The data are expressed as the means ± SEMs and are representative of two independent experiments (n = 3-5 mice/group). *\*p < 0.05 and **p < 0.01*.

Consistent with this reduced pathology, we observed a significant reduction in the frequency and number of CD8^+^ T cells in the lesion of Rag1^-/-^ mice that received CCR5^-/-^ CD8^+^ T cells compared to Rag1^-/-^ mice that received WT CD8^+^ T cells (Fig 6F-H). Additionally, Rag1^-/-^ + CCR5^-/-^ CD8 mice had a significant reduction in the frequency and number of neutrophils (CD11b^+^ Ly6G^+^ cells) (Fig 5I-K) and in neutrophils expressing pro-IL-1β (Fig 5L-N). Altogether, these results demonstrate that CD8^+^ T cells migrate to the lesion in a CCR5-dependent manner.

**Fig 6.**
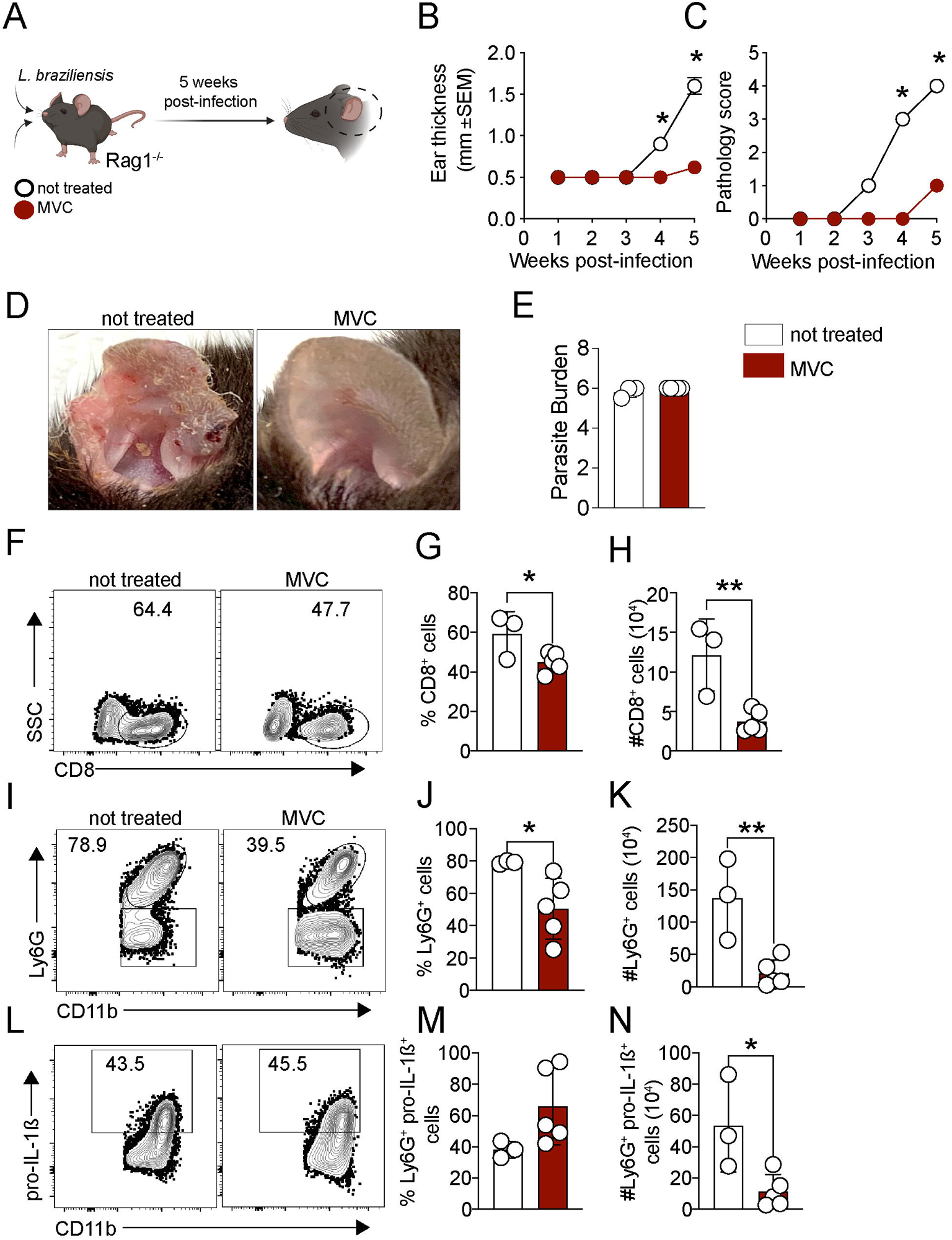
CCR5 inhibition prevents CD8^+^ T cell-dependent immunopathology. (A) Rag1^-/-^ mice were infected with *L. braziliensis*, reconstituted with CD8^+^ T cells and treated daily with maraviroc (MVC). (B) Ear thickness and (C) pathology score was evaluated weekly. (D) Representative pictures of lesions from infected-Rag1^-/-^ treated with MVC or not treated at 5 weeks post-infection. (E-N) At 5 weeks post-infection, mice were euthanized, and the ears were digested for (E) parasite quantification by limiting dilution and (F-N) flow cytometry analysis. (F) Representative dot plots and bar graphs of the (G) frequency and (H) number of CD8^+^ T cells. (I-N) The neutrophil number (CD11b^+^ Ly6G^+^ cells) and pro-IL-1β expression in the lesion were determined directly *ex vivo* by flow cytometry. (I-K) Representative dot plots and graph bars of neutrophils and (L-N) pro-IL-1β expression. The data are expressed as the means ± SEMs and are representative of two independent experiments (n = 3-5 mice). *\*p < 0.05 and **p < 0.01*.

### CCR5 inhibition by maraviroc impairs CL pathology

To investigate the therapeutic potential of blocking CCR5, we took advantage of maraviroc (MVC), an FDA-approved drug used to treat HIV infection that selectively inhibits CCR5 [27]. We tested if MVC treatment abrogated CD8^+^ T cell-mediated pathology. Infected-Rag1^-/-^ mice reconstituted with CD8^+^ T cells were treated daily with MVC by intraperitoneal injection or not treated as the control group (Fig 6A). As expected, control mice developed uncontrolled lesions with increased pathology, while mice treated with MVC showed a significant reduction in lesion size (Fig 6B) and minimum pathology (Fig 6C and 6D). No differences were observed in parasite numbers between MVC-treated and untreated mice (Fig 6E). We found that mice treated with MVC had a reduced frequency (Fig 6F and 6G) and number of CD8^+^ T cells in the lesion (Fig 6H). Additionally, MVC-treated mice had a significantly reduced frequency and number of neutrophils (CD11b+ Ly6C^+^ cells) (Fig 6I-K) and a reduced number of neutrophils expressing pro-IL-1b (Fig 6L-N). No differences were observed in macrophages, monocytes, and dendritic cell populations. Altogether, these data demonstrate that the inhibition of CCR5 prevents the CD8^+^ cells mediated pathology without affecting parasite number.

## DISCUSSION

CD8^+^ T cells mediate the destructive inflammation in *L. braziliensis* patients due to their cytolytic activity leading to cell death, NLRP3 activation, IL-1b secretion, and consequently exacerbated inflammation [7–9]. We previously demonstrated that blocking this pathway ameliorates the severity of the disease in murine models [10]. Here, we investigated the factors involved in CD8^+^ T cell migration to lesions in order to identify approaches to lessen disease severity. We identified CCR5 and its ligands, CCL3 and CCL4, as critical for CD8^+^ T cell migration to the lesion. Importantly, we demonstrated that the treatment with maraviroc, a selective inhibitor of CCR5, significantly blocked CD8^+^ T cell recruitment to leishmanial lesions and the development of severe disease without affecting parasite control.

Among the chemokine receptors related to T-cell migration analyzed in our study, we identified CCR5 as the most highly expressed in patients’ lesions and found a correlation between CCR5 and cytolytic genes, which were previously described to be associated with treatment failure [7]. CCR5 is expressed by monocytes [28], dendritic cells [29], NK cells [30,31], and lymphocytes [32,33] and regulates the trafficking and effector function of those cells. CCR5 contributes to the pathogenesis of numerous diseases, including graft-versus-host disease [34], autoimmune diseases [35],[36], and infectious diseases [13], [37]. CCR5 also promotes the migration of memory and effector-specific CD8^+^ T cells to peripheral tissues and plays a beneficial role in controlling viral and toxoplasma replication in the lungs [38] and intestine [39], respectively. However, consistent with our results, CCR5 is deleterious in situations where the cytolytic activity of CD8^+^ T cells leads to tissue damage, for example, in cerebral malaria [37], *T. cruzi*-elicited cardiomyopathy [40], acute hepatitis caused by HAV infection [13], and alopecia areata [41]. In these situations, CCR5 inhibition or deletion limits CD8^+^ T cell-mediated pathology.

The biological effect of CCR5 is mediated by its interaction with the chemokines CCL3 and CCL4 [42,43]. Our transcriptional analysis of lesions demonstrated that *L. braziliensis* patients can be classified by the high and low expression of *CCL3/4* at the lesions. Importantly, our data demonstrated that patients with *CCL3/4*^high^ expression had delayed healing of lesions and enrichment of cytolytic genes and *CCR5*. These results suggest that the increased expression of *CCL3* and *CCL4* leads to the recruitment of cytolytic CD8^+^ T cells. CCL3 and CCL4 are produced by dendritic cells[44], neutrophils [45], lymphocytes [46], and non-hematopoietic cells, such as endothelial and epithelial cells [47] in the peripheral tissue in situations of inflammation or infection, and orchestrate the immune responses by promoting the recruitment of CCR5-expressing leukocytes and also by contributing to their effector functions [45,48] [13]. Additionally, in silico tumor simulations demonstrated that CCL3 and CCL4 attract CTLs into the tumor, and the newly arriving CTLs amplify chemokine production and promote a positive feedback loop of recruitment[49]. Thus, in addition to directly promoting pathology by cell death of infected cells, it is possible that recruited cytotoxic CD8^+^ T cells at the *L. braziliensis* lesions may amplify and sustain the loop of inflammation that leads to increased pathology.

Interestingly, there is a polymorphism in CCR5 (CCR5Δ32 allele) that results in a non-functional CCR5, which is reported to prevent cell invasion by HIV-1 [50,51] and is implicated in defective cell chemotaxis [52]. A study investigating whether the CCR5 polymorphism influences the progression of cutaneous to mucocutaneous in a Brazilian cohort of *L. braziliensis-*infected patients found that no mucocutaneous patients were CCR5 polymorphism carriers, although due to the small sample size, no significant differences in the CCR5Δ32 frequency between cutaneous and mucocutaneous were found [53]. However, given the role we have described for CCR5, additional studies would be appropriate to evaluate if patients who rapidly cure *L. braziliensis* lesions are CCR5 polymorphism carriers versus patients who have a delay in healing.

To directly test the role of CCR5 expression in cytolytic CD8^+^ T cells, we tested the role of CCR5 deficient CD8 T cells in promoting increased disease Rag1^-/-^ mice. In this model, CD8^+^ T cells promote pathology by a mechanism dependent on the cytolytic function of perforin and granzyme B, which leads to inflammatory infiltration and IL-1b release, similar to the phenotype we observe in patients’ lesions [8,24]. An advantage of this model is that it allowed us to study the role of CCR5 specifically on CD8^+^ T cells, which is important since many different cells express CCR5 [28,30,32,33]. We found that CD8^+^ T cells express CCR5 preferentially at the lesions compared to the dLN. Consistent with our observation, CCR5 is mainly expressed by effector and memory CD8^+^ T cells [39],[13,37,38,41], while naïve CD8^+^ T cells express CCR5 in a transient manner in the draining lymph node [54,55]. Also, our results show that Rag1^-/-^ mice reconstituted with CD8^+^ T cells have an enrichment of *CCR5* and its ligands compared to Rag1^-/-^ mice reconstituted with CD8^+^ and CD4^+^ T cells, suggesting that this chemotactic pathway is a feature of increased pathology derived from cytolytic CD8^+^ T cells activity.

Although our results show that CCR5 has a critical role in driving CD8^+^ T cell migration to the leishmanial lesions, we cannot exclude the participation of other chemokine receptors acting to promote CD8^+^ T cell migration to the lesion. The transcriptional analysis identified enrichment of CXCR3 and its ligands CXCL9/10/11, in *L. braziliensis*-lesions compared to healthy skin. CXCR3 is required for memory CD8^+^ T cell recruitment to the lung during *Mycobacterium tuberculosis* infection [56] and during intracellular parasitic infections[37,57]. For example, CXCR3 promotes the migration of CD8^+^ T cells to the cardiac tissue and the brain in the context of *T. cruzi* and *Plasmodium* infection, respectively [37,57]. However, CXCR3 plays a beneficial role during cutaneous leishmaniasis by promoting the recruitment of Th1 cells to the lesions. Thus, *L. major*-infected CXCR3^-/-^ mice are more susceptible to infection due to a reduced number of CD4^+^T cells producing IFNγ in the lesion. Consequently, these mice fail to control parasite replication [58]. For these reasons, CXCR3 inhibition is not a good approach for host-directed therapy due to predictable adverse side effects.

Investigating the factors involved in CD8^+^ T cells migration to lesions, we found that IL-15 upregulates CCR5 expression on circulating CD8^+^ T cells from *L. braziliensis*-infected patients. IL-15 promotes the activation, proliferation, and cytotoxicity of effector and memory CD8^+^ T cells [59–61]. We previously found that IL-15 is enriched in *L. braziliensis* lesions, and inhibition of IL-15 signaling by tofacitinib ameliorates pathology in mice by dampening the cytotoxicity function of CD8^+^ T cells [6]. In the current study, we found that *IL15* is enriched in patients in the blood, reinforcing the relevance of IL-15 expression in the disease. In agreement with our data, it has been reported that IL-15 is over-produced systemically during infections [59,62,63] and inflammatory diseases [64–66], and we hypothesize that the systemic expression of IL-15 may contribute to the activation and migration of circulating CD8^+^ T cells. This idea is supported by the fact that IL-15-treated memory CD8^+^ T cells migrate from the circulation to peripheral tissues in a CCR5-dependent manner [13]. We found a positive correlation between the systemic expression of *IL15* and *CCR5* and demonstrated that IL-15 upregulates CCR5 on circulating CD8^+^ T cells. We suggest that in cutaneous leishmaniasis, systemic IL-15 may contribute to the activation of circulating CD8^+^ T cells and promote CCR5-dependent migration to the site of infection. Concomitantly, CCR5 ligands are over-expressed in leishmanial lesions, which contributes to the recruitment of CCR5-expressing CD8^+^ T cells to the site of infection. Once in the lesion, IL-15 can enhance the cytolytic activity of CD8^+^ T cells and could also contribute to the retention of CD8^+^ T cells by maintaining CCR5 expression.

The current first-line treatment of cutaneous leishmaniasis in Brazil is pentavalent antimony, which has significant side effects and often requires multiple rounds of treatments [3,4,67]. Therefore, complementing drug treatment with host-directed therapy aimed at reducing pathologic immune responses would be beneficial. Our data demonstrated the efficacy of MVC treatment in blocking the pathology induced by CD8^+^ T cells in a murine model, revealing a new possibility of effective host-directed therapy. Because MVC is an FDA–approved drug, our findings raise the possibility that it could be repurposed for treating a subset of patients with cutaneous leishmaniasis. MVC is a selective CCR5 antagonist and has been used successfully in many other situations. For example, MVC is a treatment for CCR5-tropic HIV infection [27,68,69]. MVC effectively protects against graft-versus-host disease by blocking the recruitment of alloreactive donor T-cell responses[12,70–72] and improves lesions in a murine model of alopecia areata by impairing CD8^+^ T cell migration [41]. Thus, MVC could be readily tested in cutaneous leishmaniasis patients, given its oral route of administration and known safety profile. Additionally, it was reported that *L. major*-infected CCR5^-/-^ mice developed smaller lesions and had reduced parasite number due to deficient recruitment of CD4^+^ T regulatory cells to the lesion, leading to an enhanced Th1 response[73]. Similarly, *L. donovani*-infected CCR5^-/-^ or CCL3^-/-^ mice exhibited enhanced IFNg antigen-specific in the chronic phase of the disease [74]. Therefore, we suggest that the inhibition of CCR5 might prevent the migration of pathogenic CD8^+^ T cells without interfering with the protective immune mechanisms related to the control of parasite replication. Together, these findings identified an approach that can be employed as a treatment in combination with anti-parasitic drugs to ameliorate cutaneous leishmaniasis severity.

## ACKNOWLEDGMENTS

We thank the staff at Corte de Pedra, Salvador, Bahia, Brazil for assistance with patient screening and sample collection in this study, and Drs. Helton Santiago and Liliane Monteiro Cunha for assistance. Support for this work was provided by NIH grants RO1-AI-150606 (PS) RO1-AI-143790 (PS), RO1-AI-149456 & RO1-AI-136862 (PS and LPC), P50-AI-030639 (EMC), R01-AI-162711 (FON) and RO1-AI-136862 (LPC).

## AUTHOR CONTRIBUTIONS

Conceptualization: PS, LAS; Methodology: LAS, FON, CFA, CGL; Software: CFA, DB; Validation: PS, LAS, CFA; Formal Analysis: LAS, PS, CFA, DB; Investigation: PS, LAS; Resources: PS, LPC, EMC; Data Curation: CFA, DB; Writing - Original Draft: LAS, PS; Writing - Review & Editing: PS, LAS; Visualization: LAS; Project administration: PS, EMC; Experimental work: LAS, CFA, CGL, FON; Funding Acquisition: PS, LPC, EMC; and Supervision: PS.

## METHODS

### Patients

All cutaneous leishmaniasis patients were seen at the health post in Corte de Pedra, Bahia, Brazil. The criteria for diagnosis were a clinical characteristic of cutaneous leishmaniasis and parasite confirmation by PCR or positive delayed-type hypersensitivity response to *Leishmania* antigen. This study was conducted according to the principles specified in the Declaration of Helsinki and under local ethical guidelines (Ethical Committee of the Faculdade de Medicina da Bahia, Universidade Federal da Bahia, Salvador, Bahia, Brazil, and the University of Pennsylvania Institutional Review Board), and all patients signed informed consent before enrollment into the study.

### Mice

Six-to 8-week-old female mice were used for the infection experiments. C57BL/6 mice (RRID:MGI:2159769) were purchased from Charles River, and CCR5^-/-^ (B6.129P2-Ccr5tm1Kuz/J, RRID:IMSR_JAX:005427) and Rag1^-/-^ (B6.129S7-Rag1tm1Mom/J, RRID:IMSR_JAX:002216) were purchased from the Jackson Laboratory. All mice were maintained in a specific pathogen–free facility with free access to food and water, nesting material, and housed at a temperature of 21°C at the University of Pennsylvania Animal Care Facilities. All animals were used in accordance with the recommendations in the Guide for the Care and Use of Laboratory Animals of the National Institutes of Health, and the guidelines of the University of Pennsylvania Institutional Animal Use and Care Committee. The protocol was approved by the Institutional Animal Care and Use Committee, University of Pennsylvania Animal Welfare Assurance Number.

### Parasites and bacterial cultures

*L. major* parasites (strain WHO/MHOM/IL/80/Friedlin) and *L. braziliensis* parasites (strain MHOM/BR/01/BA788) were grown in Schneider’s insect medium (GIBCO) supplemented with 20% heat-inactivated FBS, 2 mM glutamine, 100 U/mL penicillin, and 100 mg/mL streptomycin per mL. Parasitic virulence was maintained by serial passaging in BALB/c mice and by culturing *in vitro* for no more than five passages. An isolate of *S. xylosus* was cultured from the ears of *L. major* infected mice[26]. For topical associations and infections, the bacteria were cultured in Brain heart infusion (BHI) media (Remel, Lenexa, KS, USA) shaking for 12 hr at 37°C.

### Peripheral blood mononuclear cell cultures

Peripheral blood mononuclear cells (PBMCs) were isolated by centrifugation using a Ficoll-Paque Plus gradient (GE Healthcare, Cat #17-1440-02), then washed by centrifugation, and resuspended in RPMI 1640 media (Gibco) supplemented with 10% fetal bovine serum (FBS) (Gibco) and penicillin and streptomycin (Gibco). The PBMCs were adjusted to a concentration of 1 × 10^6^ cells/mL in 500 mL of RPMI 1640 media. Culturing was performed in the presence or absence of recombinant IL-15 (10 ng/mL) (Petrotech, Cat. #200-01B) and incubated for 18h at 37°C under 5% CO_2_ for 18h.

### Human cutaneous leishmaniasis transcriptional profiling by RNA-seq

The analyses carried out in this study with the lesion and blood human datasets Amorim et al. 2019, Amorim et al. 2023, and Amorim et al. 2021[7,23,24] were performed from the filtered, normalized gene expression matrix available for download as a text file on NCBI GEO accession #PRJNA682985 and #PRJNA525604. The list of six chemokine receptors associated with T-cell migration was collected based on literature[12–19]. Differential gene expression analysis was performed with edgeR R package (RRID:SCR_012802)[75]. Volcano plot was performed using DataGraph (Visual Data Tools). MCP-counter[22] and immundeconv R packages[76] were combined to estimate cytotoxic T cell abundances (CTL score) from the unstructured lesional RNA-seq dataset. *S. aureus* transcript identification, quantification, and patient stratification between *S. aureus* high vs. low were described[23]. Briefly, the dual RNA-seq computational approach was performed, in which the reads were mapped to both the human transcriptome and *S. aureus* pan-genome (#PRJNA885131) to estimate total bacterial transcript abundances.

### Intradermal infections and bacterial topical association

Infective-stage promastigotes (metacyclics) were isolated from 4–5-day old stationary culture by density gradient separation by Ficoll (Sigma-Aldrich, Cat #F9378). Mice were infected with 10^5^ *L. braziliensis* metacyclic enriched promastigotes intradermally in the left ear. Lesion development was evaluated weekly by ear thickness with a digital caliper (Fisher Scientific), and the pathology was scored from 0 to 5 based on the criteria: 0) absence of lesion; 1) swelling/redness; 2) deformation of ear pinna; 3) ulceration; 4) partial tissue loss; 5) total tissue loss. For topical associations,10^8^ CFUs of *S. xylosus* were applied to the entire mouse body using sterile cotton swabs every other day for a total of five times.

### Cell purification and adoptive transfer

Spleen cells from C57BL/6 mice were collected, erythrocytes lysed with ACK lysing buffer (Quality Biological, Cat #118-156-101), and CD8^+^ T cells were purified using a magnetic bead separation kit (Miltenyi Biotec, Cat #130-104-075). Rag1^-/-^ mice were reconstituted with CD8^+^ T cells by intravenous route (3 x 10^6^ cells/mouse) and subsequently were infected with *L. braziliensis*. Mice received 250 mg of antibody anti-CD4, clone GK1.5 (BioXCell, Cat #BE0003-1, RRID:AB_1107636) by intraperitoneal injections twice a week in the first two weeks post-infection[8].

### Ear preparation and parasite titration

Infected ears were collected, the dorsal and ventral layers of the ear were split mechanically and placed dermis side down in a 24 wells plate with 500 μl/well of RPMI with 250 mg/mL of Libarase (Roche, Cat #05401054001) and 10 mg/mL of DNase I (Sigma-Aldrich, Cat #4536282001) for 90 min at 37 °C, 5% CO_2_. The enzyme reaction was stopped with 1 mL of RPMI supplemented with 10% FBS. Ears were dissociated using a cell strainer (40mm) in PBS containing 0.05% BSA and 20 mM EDTA (Invitrogen, Cat #130-104-075), and the cell suspension was used for parasite titration. The homogenate was serially diluted (1:10) in 96-well plates and incubated at 26 °C. The number of viable parasites was calculated from the highest dilution at which parasites were observed after 7 days.

### Mouse cutaneous leishmaniasis transcriptional profiling by RNA-seq

Two RNA-seq datasets were processed for this study: Dataset 1) C57BL/6 mice were topically colonized with 10^8^ *S. xylosus* every day for a total of five applications and then intradermally infected with 2 x 10^6^ *L. major* parasites. Mice were colonized once per week for the duration of the experiment. At 5 weeks post-infection, lesions were obtained for bulk RNA-seq. Dataset 2) Rag1^-/-^ mice were infected with *L. braziliensis* and subsequently reconstituted with CD8^+^ T cells alone or CD8^+^ and CD4^+^ T cells or did not receive any T cells (φ). 5 weeks post-infection, mice were euthanized, and ears were collected in RNAlater (ThermoFisher, Cat #AM7020) for bulk RNA-seq studies. For both datasets, RNA was extracted using the RNeasy Plus Mini Kit (QIAGEN, Cat #74106) according to the manufacturer’s instructions and used to prepare Poly(A)+-enriched cDNA libraries Illumina TruSeq Stranded mRNA library prep workflow. Ribo-Zero Gold rRNA depletion (Illumina, Cat # MRZG12324) was used to remove ribosomal content. Quality assessment and quantification of RNA preparations and libraries were carried out using an Agilent 4200 TapeStation and Qubit 3, respectively. Samples were sequenced on an Illumina NextSeq 500 The raw reads were mapped to the mouse reference transcriptome (Ensembl; Mus musculus) using Kallisto[77]. All subsequent analyses were conducted using the statistical computing environment R, RStudio, and Bioconductor. Transcript quantification, normalization, filtering for highly expressed genes, variance-stabilization, and Differential Gene Expression analysis were performed as described previously [23]. Raw sequence data are available at the NCBI GEO BioProjects XXXX and XXXX.

### Flow cytometry

Single-cell suspensions were stained with LIVE/DEAD Fixable Aqua Dead Cell Stain Kit (Molecular Probes, Cat #L34957) and subsequently incubated with anti-CD16/CD32 (eBioscience, Cat #14-0161-86, clone 93) and 10% rat-IgG1 (Sigma-Aldrich, Cat #I8015) in PBS containing 0.1% BSA (Sigma-Aldrich). For surface staining, cells were incubated with monoclonal antibodies anti-mouse (anti-CD45 APCcy7 [clone30-F11, Cat #103116, RRID:AB_312981], anti-CD3 BV605 [clone 172A, Biolegend, Cat #100237, RRID:AB_2562039], anti-CD8 BV711 [clone 53-6.7, Biolegend, Cat #100759, RRID:AB_2563510], anti-Ly6G pacific blue [clone 1A8, Biolegend, Cat #127612, RRID:AB_2251161], anti-F4/80 BV605 [clone BM8, Biolegend, Cat #123133, RRID:AB_2562305], anti-CD11c BV711 [clone N418, Biolegend, Cat #117349, RRID:AB_2563905], anti-Ly6C BV785 [clone HK1.4, Biolegend, Cat #128041, RRID:AB_2565852], anti-MHCII Alexa fluor-700 [clone M5/114.15.2, eBiosciences, Cat #56-5321-82, RRID:AB_494009], anti-CD44 eF450 [clone IM7, Biosciences, Cat #48-0441-82, RRID:AB_1272246], and anti-CCR5 PE [clone C34-3448, eBiosciences, Cat #559923, RRID:AB_397377] or anti-human antibodies (anti-CD45 Pecy7 [clone HI30, eBiosciences, Cat #25-0459-42, RRID:AB_1944375], anti-CD8 Percp-cy5.5 [clone SK1, Biolegend, Cat #344710, RRID:AB_2044010], anti-TCRa/b APC [clone IP26, eBiosciences, Cat #17-9986-42, RRID:AB_10597896] and anti-CCR5 BV421 [clone 2D7, BD Biosciences, Cat #562576, RRID:AB_2737661]), followed by fixation with 2% of formaldehyde and permeablization with 0.2% saponin/PBS. For intracellular staining, cells obtained from mice were permeabilized with 0.4% saponin buffer and stained *ex vivo* for pro-IL-1b APC (clone NJTEN3, eBiosciences, Cat #17-7114-80, RRID:AB_10670739). Data were collected using LSRIII Fortessa (BD Biosciences) and analyzed using FlowJo software (Tree Star, RRID:SCR_008520).

### Maraviroc treatment

Mice were intraperitoneally treated with 20 mg/kg of Maraviroc (Cayman Chemical, Cat #14641, CAS Number 376348-65-1) diluted in 10% DMSO (Sigma-Aldrich, Cat # D8418, CAS number 67-68-5) (200 ml/mouse) daily.

#### Quantification and statistical analysis

Data are shown as means ± SEM. For mouse and human experiments, statistical significance was determined using the two-tailed unpaired Student’s t-test or one-way ANOVA. Pearson correlation coefficient was used to determine the correlation between log2 expressions of genes from human skin and peripheral blood transcripts. All statistical analysis was calculated using GraphPad Prism version 10 (GraphPad Software, RRID:SCR_002798). Differences were considered significant when *p < 0.05, **p ≤ 0.01, ***p ≤ 0.001, ****p < .0001. For human and mouse studies, specific sample sizes are represented by n and are indicated in figure legends.

### Data and code availability

RNA-seq data and clinical metadata from patients’ lesions and blood in this study are derived from published transcriptional profiling and are available on NCBI GEO accession #PRJNA682985, #PRJNA525604, and #PRJNA885131. All original code has been deposited at GEO and is publicly available as of the date of publication. The complete RNA-seq data analysis, R code, file inputs, and outputs used to perform the transcriptional analysis presented in this current manuscript are available on the GitHub repository “CCR5_LAS” (https://github.com/camilafarias112/CCR5_LAS). DOIs are listed in the key resources table.

## References

1. Lago AS do, Nascimento M, Carvalho AM, Lago N, Silva J, Queiroz JR, et al. The elderly respond to antimony therapy for cutaneous leishmaniasis similarly to young patients but have severe adverse reactions. Am J Trop Med Hyg. 2018;98: 1317–1324. doi:10.4269/ajtmh.17-0736

2. Ponte-Sucre A, Gamarro F, Dujardin J-C, Barrett MP, López-Vélez R, García-Hernández R, et al. Drug resistance and treatment failure in leishmaniasis: A 21st century challenge. PLoS Negl Trop Dis. 2017;11: e0006052. doi:10.1371/journal.pntd.0006052

3. Oliveira-Neto MP, Schubach A, Mattos M, Goncalves-Costa SC, Pirmez C. A low-dose antimony treatment in 159 patients with American cutaneous leishmaniasis: extensive follow-up studies (up to 10 years). Am J Trop Med Hyg. 1997;57: 651–655. doi:10.4269/ajtmh.1997.57.651

4. Arevalo J, Ramirez L, Adaui V, Zimic M, Tulliano G, Miranda-Verástegui C, et al. Influence of Leishmania (Viannia) species on the response to antimonial treatment in patients with American tegumentary leishmaniasis. J Infect Dis. 2007;195: 1846–1851. doi:10.1086/518041

5. Costa RS, Carvalho LP, Campos TM, Magalhães AS, Passos ST, Schriefer A, et al. Early Cutaneous Leishmaniasis Patients Infected With Leishmania braziliensis Express Increased Inflammatory Responses After Antimony Therapy. J Infect Dis. 2018;217: 840–850. doi:10.1093/infdis/jix627

6. Novais FO, Nguyen BT, Scott P. Granzyme B inhibition by tofacitinib blocks the pathology induced by CD8 T cells in cutaneous leishmaniasis. J Invest Dermatol. 2021;141: 575–585. doi:10.1016/j.jid.2020.07.011

7. Amorim CF, Novais FO, Nguyen BT, Misic AM, Carvalho LP, Carvalho EM, et al. Variable gene expression and parasite load predict treatment outcome in cutaneous leishmaniasis. Sci Transl Med. 2019;11. doi:10.1126/scitranslmed.aax4204

8. Novais FO, Carvalho LP, Graff JW, Beiting DP, Ruthel G, Roos DS, et al. Cytotoxic T cells mediate pathology and metastasis in cutaneous leishmaniasis. PLoS Pathog. 2013;9: e1003504. doi:10.1371/journal.ppat.1003504

9. Novais FO, Carvalho LP, Passos S, Roos DS, Carvalho EM, Scott P, et al. Genomic profiling of human Leishmania braziliensis lesions identifies transcriptional modules associated with cutaneous immunopathology. J Invest Dermatol. 2015;135: 94–101. doi:10.1038/jid.2014.305

10. Novais FO, Carvalho AM, Clark ML, Carvalho LP, Beiting DP, Brodsky IE, et al. CD8+ T cell cytotoxicity mediates pathology in the skin by inflammasome activation and IL-1β production. PLoS Pathog. 2017;13: e1006196. doi:10.1371/journal.ppat.1006196

11. Carvalho AM, Novais FO, Paixão CS, de Oliveira CI, Machado PRL, Carvalho LP, et al. Glyburide, a NLRP3 Inhibitor, Decreases Inflammatory Response and Is a Candidate to Reduce Pathology in Leishmania braziliensis Infection. J Invest Dermatol. 2020;140: 246–249.e2. doi:10.1016/j.jid.2019.05.025

12. Murai M, Yoneyama H, Harada A, Yi Z, Vestergaard C, Guo B, et al. Active participation of CCR5(+)CD8(+) T lymphocytes in the pathogenesis of liver injury in graft-versus-host disease. J Clin Invest. 1999;104: 49–57. doi:10.1172/JCI6642

13. Seo I-H, Eun HS, Kim JK, Lee H, Jeong S, Choi SJ, et al. IL-15 enhances CCR5-mediated migration of memory CD8+ T cells by upregulating CCR5 expression in the absence of TCR stimulation. Cell Rep. 2021;36: 109438. doi:10.1016/j.celrep.2021.109438

14. Kurachi M, Kurachi J, Suenaga F, Tsukui T, Abe J, Ueha S, et al. Chemokine receptor CXCR3 facilitates CD8(+) T cell differentiation into short-lived effector cells leading to memory degeneration. J Exp Med. 2011;208: 1605–1620. doi:10.1084/jem.20102101

15. Hu JK, Kagari T, Clingan JM, Matloubian M. Expression of chemokine receptor CXCR3 on T cells affects the balance between effector and memory CD8 T-cell generation. Proc Natl Acad Sci USA. 2011;108: E118–27. doi:10.1073/pnas.1101881108

16. Wein AN, McMaster SR, Takamura S, Dunbar PR, Cartwright EK, Hayward SL, et al. CXCR6 regulates localization of tissue-resident memory CD8 T cells to the airways. J Exp Med. 2019;216: 2748–2762. doi:10.1084/jem.20181308

17. Imai T, Nagira M, Takagi S, Kakizaki M, Nishimura M, Wang J, et al. Selective recruitment of CCR4-bearing Th2 cells toward antigen-presenting cells by the CC chemokines thymus and activation-regulated chemokine and macrophage-derived chemokine. Int Immunol. 1999;11: 81–88. doi:10.1093/intimm/11.1.81

18. Zingoni A, Soto H, Hedrick JA, Stoppacciaro A, Storlazzi CT, Sinigaglia F, et al. The chemokine receptor CCR8 is preferentially expressed in Th2 but not Th1 cells. J Immunol. 1998;161: 547– 551.

19. Valentine KM, Hoyer KK. CXCR5+ CD8 T cells: protective or pathogenic? Front Immunol. 2019;10: 1322. doi:10.3389/fimmu.2019.01322

20. Faria DR, Souza PEA, Durães FV, Carvalho EM, Gollob KJ, Machado PR, et al. Recruitment of CD8(+) T cells expressing granzyme A is associated with lesion progression in human cutaneous leishmaniasis. Parasite Immunol. 2009;31: 432–439. doi:10.1111/j.1365-3024.2009.01125.x

21. Santos C da S, Boaventura V, Ribeiro Cardoso C, Tavares N, Lordelo MJ, Noronha A, et al. CD8(+) granzyme B(+)-mediated tissue injury vs. CD4(+)IFNγ(+)-mediated parasite killing in human cutaneous leishmaniasis. J Invest Dermatol. 2013;133: 1533–1540. doi:10.1038/jid.2013.4

22. Becht E, Giraldo NA, Lacroix L, Buttard B, Elarouci N, Petitprez F, et al. Estimating the population abundance of tissue-infiltrating immune and stromal cell populations using gene expression. Genome Biol. 2016;17: 218. doi:10.1186/s13059-016-1070-5

23. Farias Amorim C, Lovins VM, Singh TP, Novais FO, Harris JC, Lago AS, et al. The skin microbiome enhances disease through IL-1b and delays healing in cutaneous leishmaniasis patients. medRxiv. 2023. doi:10.1101/2023.02.02.23285247

24. Farias Amorim C, O Novais F, Nguyen BT, Nascimento MT, Lago J, Lago AS, et al. Localized skin inflammation during cutaneous leishmaniasis drives a chronic, systemic IFN-γ signature. PLoS Negl Trop Dis. 2021;15: e0009321. doi:10.1371/journal.pntd.0009321

25. Sacramento LA, Farias Amorim C, Campos TM, Saldanha M, Arruda S, Carvalho LP, et al. NKG2D promotes CD8 T cell-mediated cytotoxicity and is associated with treatment failure in human cutaneous leishmaniasis. PLoS Negl Trop Dis. 2023;17: e0011552. doi:10.1371/journal.pntd.0011552

26. Gimblet C, Meisel JS, Loesche MA, Cole SD, Horwinski J, Novais FO, et al. Cutaneous Leishmaniasis Induces a Transmissible Dysbiotic Skin Microbiota that Promotes Skin Inflammation. Cell Host Microbe. 2017;22: 13–24.e4. doi:10.1016/j.chom.2017.06.006

27. Tan Q, Zhu Y, Li J, Chen Z, Han GW, Kufareva I, et al. Structure of the CCR5 chemokine receptor-HIV entry inhibitor maraviroc complex. Science. 2013;341: 1387–1390. doi:10.1126/science.1241475

28. Rawat K, Tewari A, Li X, Mara AB, King WT, Gibbings SL, et al. CCL5-producing migratory dendritic cells guide CCR5+ monocytes into the draining lymph nodes. J Exp Med. 2023;220. doi:10.1084/jem.20222129

29. Aliberti J, Reis e Sousa C, Schito M, Hieny S, Wells T, Huffnagle GB, et al. CCR5 provides a signal for microbial induced production of IL-12 by CD8 alpha+ dendritic cells. Nat Immunol. 2000;1: 83–87. doi:10.1038/76957

30. Khan IA, Thomas SY, Moretto MM, Lee FS, Islam SA, Combe C, et al. CCR5 is essential for NK cell trafficking and host survival following Toxoplasma gondii infection. PLoS Pathog. 2006;2: e49. doi:10.1371/journal.ppat.0020049

31. Ajuebor MN, Wondimu Z, Hogaboam CM, Le T, Proudfoot AEI, Swain MG. CCR5 deficiency drives enhanced natural killer cell trafficking to and activation within the liver in murine T cell-mediated hepatitis. Am J Pathol. 2007;170: 1975–1988. doi:10.2353/ajpath.2007.060690

32. Kroetz DN, Deepe GS. CCR5 dictates the equilibrium of proinflammatory IL-17+ and regulatory Foxp3+ T cells in fungal infection. J Immunol. 2010;184: 5224–5231. doi:10.4049/jimmunol.1000032

33. Loetscher P, Uguccioni M, Bordoli L, Baggiolini M, Moser B, Chizzolini C, et al. CCR5 is characteristic of Th1 lymphocytes. Nature. 1998;391: 344–345. doi:10.1038/34814

34. Ichiki Y, Bowlus CL, Shimoda S, Ishibashi H, Vierling JM, Gershwin ME. T cell immunity and graft-versus-host disease (GVHD). Autoimmun Rev. 2006;5: 1–9. doi:10.1016/j.autrev.2005.02.006

35. Sellebjerg F, Madsen HO, Jensen CV, Jensen J, Garred P. CCR5 delta32, matrix metalloproteinase-9 and disease activity in multiple sclerosis. J Neuroimmunol. 2000;102: 98–106. doi:10.1016/s0165-5728(99)00166-6

36. Carvalho-Pinto C, García MI, Gómez L, Ballesteros A, Zaballos A, Flores JM, et al. Leukocyte attraction through the CCR5 receptor controls progress from insulitis to diabetes in non-obese diabetic mice. Eur J Immunol. 2004;34: 548–557. doi:10.1002/eji.200324285

37. Belnoue E, Kayibanda M, Deschemin J-C, Viguier M, Mack M, Kuziel WA, et al. CCR5 deficiency decreases susceptibility to experimental cerebral malaria. Blood. 2003;101: 4253–4259. doi:10.1182/blood-2002-05-1493

38. Kohlmeier JE, Miller SC, Smith J, Lu B, Gerard C, Cookenham T, et al. The chemokine receptor CCR5 plays a key role in the early memory CD8+ T cell response to respiratory virus infections. Immunity. 2008;29: 101–113. doi:10.1016/j.immuni.2008.05.011

39. Luangsay S, Kasper LH, Rachinel N, Minns LA, Mennechet FJD, Vandewalle A, et al. CCR5 mediates specific migration of Toxoplasma gondii-primed CD8 lymphocytes to inflammatory intestinal epithelial cells. Gastroenterology. 2003;125: 491–500. doi:10.1016/s0016-5085(03)00903-x

40. Gibaldi D, Vilar-Pereira G, Pereira IR, Silva AA, Barrios LC, Ramos IP, et al. CCL3/Macrophage Inflammatory Protein-1α Is Dually Involved in Parasite Persistence and Induction of a TNF- and IFNγ-Enriched Inflammatory Milieu in Trypanosoma cruzi-Induced Chronic Cardiomyopathy. Front Immunol. 2020;11: 306. doi:10.3389/fimmu.2020.00306

41. Ito T, Suzuki T, Funakoshi A, Fujiyama T, Tokura Y. CCR5 is a novel target for the treatment of experimental alopecia areata. J Cutan Immunol Allergy. 2020;3: 24–32. doi:10.1002/cia2.12092

42. Raport CJ, Gosling J, Schweickart VL, Gray PW, Charo IF. Molecular cloning and functional characterization of a novel human CC chemokine receptor (CCR5) for RANTES, MIP-1beta, and MIP-1alpha. J Biol Chem. 1996;271: 17161–17166. doi:10.1074/jbc.271.29.17161

43. Combadiere C, Ahuja SK, Tiffany HL, Murphy PM. Cloning and functional expression of CC CKR5, a human monocyte CC chemokine receptor selective for MIP-1(alpha), MIP-1(beta), and RANTES. J Leukoc Biol. 1996;60: 147–152. doi:10.1002/jlb.60.1.147

44. Bystry RS, Aluvihare V, Welch KA, Kallikourdis M, Betz AG. B cells and professional APCs recruit regulatory T cells via CCL4. Nat Immunol. 2001;2: 1126–1132. doi:10.1038/ni735

45. Charmoy M, Brunner-Agten S, Aebischer D, Auderset F, Launois P, Milon G, et al. Neutrophil-derived CCL3 is essential for the rapid recruitment of dendritic cells to the site of Leishmania major inoculation in resistant mice. PLoS Pathog. 2010;6: e1000755. doi:10.1371/journal.ppat.1000755

46. Cook DN, Smithies O, Strieter RM, Frelinger JA, Serody JS. CD8+ T cells are a biologically relevant source of macrophage inflammatory protein-1 alpha in vivo. J Immunol. 1999;162: 5423– 5428.

47. Maurer M, von Stebut E. Macrophage inflammatory protein-1. Int J Biochem Cell Biol. 2004;36: 1882–1886. doi:10.1016/j.biocel.2003.10.019

48. Lindell DM, Standiford TJ, Mancuso P, Leshen ZJ, Huffnagle GB. Macrophage inflammatory protein 1alpha/CCL3 is required for clearance of an acute Klebsiella pneumoniae pulmonary infection. Infect Immun. 2001;69: 6364–6369. doi:10.1128/IAI.69.10.6364-6369.2001

49. Galeano Niño JL, Pageon SV, Tay SS, Colakoglu F, Kempe D, Hywood J, et al. Cytotoxic T cells swarm by homotypic chemokine signalling. eLife. 2020;9. doi:10.7554/eLife.56554

50. Liu R, Paxton WA, Choe S, Ceradini D, Martin SR, Horuk R, et al. Homozygous defect in HIV-1 coreceptor accounts for resistance of some multiply-exposed individuals to HIV-1 infection. Cell. 1996;86: 367–377. doi:10.1016/s0092-8674(00)80110-5

51. Theodorou I, Meyer L, Magierowska M, Katlama C, Rouzioux C. HIV-1 infection in an individual homozygous for CCR5 delta 32. Seroco Study Group. Lancet. 1997;349: 1219–1220.

52. Yang X, Ahmad T, Gogus F, Verity D, Wallace GR, Madanat W, et al. Analysis of the CC chemokine receptor 5 (CCR5) Delta32 polymorphism in Behçet’s disease. Eur J Immunogenet. 2004;31: 11–14. doi:10.1111/j.1365-2370.2004.00444.x

53. Brajão de Oliveira K, Reiche EMV, Kaminami Morimoto H, Pelegrinelli Fungaro MH, Estevão D, Pontello R, et al. Analysis of the CC chemokine receptor 5 delta32 polymorphism in a Brazilian population with cutaneous leishmaniasis. J Cutan Pathol. 2007;34: 27–32. doi:10.1111/j.1600-0560.2006.00573.x

54. Castellino F, Huang AY, Altan-Bonnet G, Stoll S, Scheinecker C, Germain RN. Chemokines enhance immunity by guiding naive CD8+ T cells to sites of CD4+ T cell-dendritic cell interaction. Nature. 2006;440: 890–895. doi:10.1038/nature04651

55. Askew D, Su CA, Barkauskas DS, Dorand RD, Myers J, Liou R, et al. Transient Surface CCR5 Expression by Naive CD8+ T Cells within Inflamed Lymph Nodes Is Dependent on High Endothelial Venule Interaction and Augments Th Cell-Dependent Memory Response. J Immunol. 2016;196: 3653–3664. doi:10.4049/jimmunol.1501176

56. Kohlmeier JE, Reiley WW, Perona-Wright G, Freeman ML, Yager EJ, Connor LM, et al. Inflammatory chemokine receptors regulate CD8(+) T cell contraction and memory generation following infection. J Exp Med. 2011;208: 1621–1634. doi:10.1084/jem.20102110

57. Pontes Ferreira C, Moro Cariste L, Henrique Noronha I, Fernandes Durso D, Lannes-Vieira J, Ramalho Bortoluci K, et al. CXCR3 chemokine receptor contributes to specific CD8+ T cell activation by pDC during infection with intracellular pathogens. PLoS Negl Trop Dis. 2020;14: e0008414. doi:10.1371/journal.pntd.0008414

58. Rosas LE, Barbi J, Lu B, Fujiwara Y, Gerard C, Sanders VM, et al. CXCR3-/- mice mount an efficient Th1 response but fail to control Leishmania major infection. Eur J Immunol. 2005;35: 515–523. doi:10.1002/eji.200425422

59. Kim J, Chang D-Y, Lee HW, Lee H, Kim JH, Sung PS, et al. Innate-like Cytotoxic Function of Bystander-Activated CD8+ T Cells Is Associated with Liver Injury in Acute Hepatitis A. Immunity. 2018;48: 161–173.e5. doi:10.1016/j.immuni.2017.11.025

60. Sowell RT, Goldufsky JW, Rogozinska M, Quiles Z, Cao Y, Castillo EF, et al. IL-15 Complexes Induce Migration of Resting Memory CD8 T Cells into Mucosal Tissues. J Immunol. 2017;199: 2536–2546. doi:10.4049/jimmunol.1501638

61. Younes S-A, Freeman ML, Mudd JC, Shive CL, Reynaldi A, Panigrahi S, et al. IL-15 promotes activation and expansion of CD8+ T cells in HIV-1 infection. J Clin Invest. 2016;126: 2745–2756. doi:10.1172/JCI85996

62. Frahm M, Goswami ND, Owzar K, Hecker E, Mosher A, Cadogan E, et al. Discriminating between latent and active tuberculosis with multiple biomarker responses. Tuberculosis (Edinb). 2011;91: 250–256. doi:10.1016/j.tube.2011.02.006

63. Kakumu S, Okumura A, Ishikawa T, Yano M, Enomoto A, Nishimura H, et al. Serum levels of IL-10, IL-15 and soluble tumour necrosis factor-alpha (TNF-alpha) receptors in type C chronic liver disease. Clin Exp Immunol. 1997;109: 458–463. doi:10.1046/j.1365-2249.1997.4861382.x

64. Kirman I, Nielsen OH. Increased numbers of interleukin-15-expressing cells in active ulcerative colitis. Am J Gastroenterol. 1996;91: 1789–1794.

65. Kivisäkk P, Matusevicius D, He B, Söderström M, Fredrikson S, Link H. IL-15 mRNA expression is up-regulated in blood and cerebrospinal fluid mononuclear cells in multiple sclerosis (MS). Clin Exp Immunol. 1998;111: 193–197. doi:10.1046/j.1365-2249.1998.00478.x

66. Kuczyński S, Winiarska H, Abramczyk M, Szczawińska K, Wierusz-Wysocka B, Dworacka M. IL-15 is elevated in serum patients with type 1 diabetes mellitus. Diabetes Res Clin Pract. 2005;69: 231–236. doi:10.1016/j.diabres.2005.02.007

67. Unger A, O’Neal S, Machado PRL, Guimarães LH, Morgan DJ, Schriefer A, et al. Association of treatment of American cutaneous leishmaniasis prior to ulcer development with high rate of failure in northeastern Brazil. Am J Trop Med Hyg. 2009;80: 574–579. doi:10.4269/ajtmh.2009.80.574

68. Lagane B, Garcia-Perez J, Kellenberger E. Modeling the allosteric modulation of CCR5 function by Maraviroc. Drug Discov Today Technol. 2013;10: e297–305. doi:10.1016/j.ddtec.2012.07.011

69. Garcia-Perez J, Rueda P, Alcami J, Rognan D, Arenzana-Seisdedos F, Lagane B, et al. Allosteric model of maraviroc binding to CC chemokine receptor 5 (CCR5). J Biol Chem. 2011;286: 33409– 33421. doi:10.1074/jbc.M111.279596

70. Reshef R, Luger SM, Hexner EO, Loren AW, Frey NV, Nasta SD, et al. Blockade of lymphocyte chemotaxis in visceral graft-versus-host disease. N Engl J Med. 2012;367: 135–145. doi:10.1056/NEJMoa1201248

71. Moy RH, Huffman AP, Richman LP, Crisalli L, Wang XK, Hoxie JA, et al. Clinical and immunologic impact of CCR5 blockade in graft-versus-host disease prophylaxis. Blood. 2017;129: 906–916. doi:10.1182/blood-2016-08-735076

72. Reshef R, Ganetsky A, Acosta EP, Blauser R, Crisalli L, McGraw J, et al. Extended CCR5 Blockade for Graft-versus-Host Disease Prophylaxis Improves Outcomes of Reduced-Intensity Unrelated Donor Hematopoietic Cell Transplantation: A Phase II Clinical Trial. Biol Blood Marrow Transplant. 2019;25: 515–521. doi:10.1016/j.bbmt.2018.09.034

73. Yurchenko E, Tritt M, Hay V, Shevach EM, Belkaid Y, Piccirillo CA. CCR5-dependent homing of naturally occurring CD4+ regulatory T cells to sites of Leishmania major infection favors pathogen persistence. J Exp Med. 2006;203: 2451–2460. doi:10.1084/jem.20060956

74. Sato N, Kuziel WA, Melby PC, Reddick RL, Kostecki V, Zhao W, et al. Defects in the generation of IFN-gamma are overcome to control infection with Leishmania donovani in CC chemokine receptor (CCR) 5-, macrophage inflammatory protein-1 alpha-, or CCR2-deficient mice. J Immunol. 1999;163: 5519–5525.

75. Robinson MD, McCarthy DJ, Smyth GK. edgeR: a Bioconductor package for differential expression analysis of digital gene expression data. Bioinformatics. 2010;26: 139–140. doi:10.1093/bioinformatics/btp616

76. Sturm G, Finotello F, List M. Immunedeconv: An R Package for Unified Access to Computational Methods for Estimating Immune Cell Fractions from Bulk RNA-Sequencing Data. Methods Mol Biol. 2020;2120: 223–232. doi:10.1007/978-1-0716-0327-7_16

77. Bray NL, Pimentel H, Melsted P, Pachter L. Near-optimal probabilistic RNA-seq quantification. Nat Biotechnol. 2016;34: 525–527. doi:10.1038/nbt.3519 Sciwheel inserting bibliography…Sciwheel inserting bibliography.

